# *Pulsatilla hezijianensis*, (Ranunculaceae), a new species from Beijing, China

**DOI:** 10.1101/2025.07.24.666620

**Authors:** Huan-Yu Wu, Shuang Qin, Wen-He Li, Jia-Min Xiao, Bo-Wen Liu, Jian He, Lei Xie

## Abstract

*Pulsatilla hezijianensis* H.Y. Wu & L. Xie, a new species of Ranunculaceae from Hezijian Village (Beijing, China), is described. This taxon resembles *P. sukaczevii* and *P. tenuiloba* in its finely dissected pinnate leaves but is distinguished by its smaller stature (3–15 cm tall), shorter scapes, smaller fruit heads, and sepals with white inner surfaces and pale blue outer surfaces. Molecular phylogenetic analyses based on nuclear (nrITS) and plastid markers (*rbc*L, *acc*D*-psa*I, *trn*L intron) revealed that the Hezijian population forms a distinct clade, sister to *P. tenuiloba* but with significant genetic divergence. Morphological comparisons with allied species, coupled with its distribution and phenology, support its recognition as a new species. The restricted distribution and small population size warrant its conservation status as Critically Endangered (CR) under IUCN criteria.

## Introduction

The genus *Pulsatilla* Miller (1754: without pagination) (Pasque-flowers), a moderately sized group within the buttercup family, comprises approximately 40 perennial herbaceous species distributed across temperate montane and boreal regions of the Northern Hemisphere (Aichele & Schwegleb 1957, Tamura 1995, Grey-Wilson 2014). The genus *Pulsatilla* contains many important medicinal plants. The roots and rhizomes of *Pulsatilla* species are used to treat dysentery, while some species are also effective against malaria, wounds, and other ailments. Additionally, they can be used as natural pesticides to control agricultural pests. Moreover, *Pulsatilla* species serve as early-spring ornamental flowers, highlighting their significant horticultural value (Zhao 1988). Northern Asia harbors the highest species diversity of *Pulsatilla*, encompassing approximately 30 species., and more than 11 species were reported in China (Wang & Bartholomew 2001, Zhang *et al.* 2022).

Plants of *Pulsatilla* typically grow from a stout, vertical rhizome and bear basal leaves that are deeply pinnately or palmately lobed, often covered with silky hairs. They produce solitary, bell-shaped to cup-shaped flowers on erect stems, with showy petal-like sepals (usually in shades of purple, violet, yellow, or white) and numerous stamens. After flowering, the styles elongate into feathery, plume-like appendages, forming ornamental achene heads that aid in wind dispersal (Tamura 1995). Most species bloom in early spring, with flowers appearing before or alongside the new leaves. Despite some molecular phylogenetic studies supporting the inclusion of *Pulsatilla* within *Anemone* Linnaeus (1753: 538) (Hoot *et al.* 2012), most of the current studies favor the generic rank of this group or at least prefer to maintain the genus name *Pulsatilla* in their studies (Ronikier *et al.* 2008, Jiang *et al.* 2017, Sramkó *et al.* 2019, Li *et al.* 2019, 2020).

*Pulsatilla* exhibits a highly complex pattern of morphological variation within the genus (Grey-Wilson 2014). Conventional taxonomic methods face challenges in distinguishing wild *Pulsatilla* species due to overlapping and intermediate morphological traits among many taxa. As mentioned by Li *et al.* (2019), several key morphological traits traditionally used for *Pulsatilla* classification, such as the number of lateral leaf lobe pairs, flower color, and flower nodding before anthesis, exhibit considerable intraspecific and even intrapopulational variation. Therefore, molecular phylogenetic evidence is particularly crucial for the identification and classification of *Pulsatilla* species.

In recent years, a morphologically distinctive *Pulsatilla* was discovered on the slopes of Hezijian Village in Beijing’s Changping District. Unlike *P. chinensis* (Bunge 1833: 76) Regel (1861: 5), the most common Pasque-flower in Beijing, this species is notably smaller in size and has leaves that are finely dissected multiple times.This plant blooms in April before its leaves fully unfold, featuring white inner surfaces and pale bluish-purple outer surfaces on its sepals. Although the Pasque-flower found in Changping does resemble *P. sukaczevii* Juzepczuk (1937: 301) (and was identified as *P. sukaczevii*) in certain aspects, such as small in size, blooming before leaf expansion, and having 2–3 times pinnately dissected leaves, typically with four pairs of lateral lobes, it differs markedly from the *P. sukaczevii* we observed in Inner Mongolia. The Hezijian plants are notably smaller and more delicate, with relatively smaller flowers that are white inside and pale purple outside. In contrast, *P. sukaczevii* from Inner Mongolia is somewhat larger and more robust, with yellow or yellowish-white flowers.

In order to accurately identify this species and clarify its phylogenetic position, we conducted field surveys in Hezijian Village, Changping Dist. during the spring of 2023 and the summer of 2025, collecting specimens for morphological examination and molecular phylogenetic analysis. The objectives of this study are to determine the taxonomic and phylogenetic position of this plant through integrated morphological and molecular evidences, thereby contributing to a better understanding of China’s biodiversity and supporting its conservation efforts.

## Materials and methods

### Morphological analysis

We checked specimens of *Pulsatilla* in China housed at BJFC, PE, KUN, K and BM, as well as specimen photos available in several web databases, such as China Virtual Herbarium (https://www.cvh.ac.cn), Plant Photo Bank of China (PPBC. https://ppbc.iplant.cn), website of Komarov Botanical Institute Herbarium (https://www.binran.ru/en/collections/herbarium/), and jstor (https://plants.jstor.org). We also studied the morphology of the new species and other related species using living plants growing in the field and green house of Beijing Forestry University. All morphological characters were observed under dissecting microscopes and described following the terminology used in Wang & Bartholomew (2001). Literature studies included taxonomic revisions in Eurasia and monographs (Juzepczuk 1937, Aichele & Schwegleb 1957, Zhao 1988, Tamura 1995, Grey-Wilson 2014, Wang & Bartholomew 2001), and recently published phylogenetic studies (Ronikier *et al.* 2008, Hoot *et al.* 2012, Jiang *et al.* 2017, Sramkó *et al.* 2019, Li *et al.* 2019, 2020, Xiang *et al.* 2025).

### Plant sampling and choosing molecular markers

In this study, we conducted field surveys and collected leaf samples of the new specie*s* in Changping Dist., Beijing, China. Multiple individuals were collected from field, and three of them were used for phylogenetic reconstruction. Since the study by Sramkó *et al.* (2019) represents the most comprehensive phylogenetic analysis of *Pulsatilla* to date, encompassing the widest taxonomic sampling within the genus, we adopted their dataset as the foundational framework for our phylogenetic reconstruction. In this study, we included other 31 species (49 samples) of *Pulsatilla* in our molecular phylogenetic analysis (Table S1). They were all sourced from Sramkó *et al.* (2019). We excluded two *Pulsatilla* samples with extensive missing data from their study in our analysis. Five samples of *Anemone* and *Hepatica* Miller (1754: without pagination) were chosen as the outgroups.

For the three newly collected samples, we obtained genome skimming data through next-generation sequencing method. Using the DNA regions in Sramkó *et al.* (2019) as the references, we assembled these target DNA regions from the genome skimming data using Geneious v.Prime (Kearse *et al.* 2012). Sramkó *et al.* (2019) used two nuclear (nrITS and MLH1) and three plastid regions (*rbc*L, *acc*D-*psa*I, *trn*L intron) for their phylogenetic analysis. Because MLH1 is a low copy nuclear gene and our genome skimming data failed to assemble it, we only use nrITS and three plastid markers in this study. Voucher specimens of the newly generated molecular data were deposited in the Herbarium of Beijing Forestry University (BJFC, herbarium code follows Thiers 2017).

### Genome skimming sequencing

Genomic DNA was extracted from silica-dried leaves using a commercial DNA isolation kit (Tiangen Biotech, Beijing, China) in accordance with the manufacturer’s instructions. The extraction was conducted at Berry Genomics Co., Ltd. (https://www.bioon.com.cn/company/index/df7a2b444074). For library construction, 1 μg of DNA from each sample was prepared using the VAHTS Universal DNA Library Prep Kit for MGI (Vazyme, Nanjing, China). The libraries were then sequenced in a 2×150 bp paired-end format on the DNBSEQ-T7 platform (BGI, Shenzhen, China), yielding approximately 6 Gbp of raw data per sample.

### Phylogenetic analyses

After assembling the four DNA regions from the newly sequenced species, sequences of each region were aligned using MAFFT v.6.833 (Katoh *et al.* 2005) with iterative manual optimization in Geneious v.Prime (Kearse *et al.* 2012), followed by pruning of ambiguous alignment regions according to Sramkó *et al.* (2019).

Because the separate regions provide limited phylogenetic information, phylogenetic analysis is conducted on the concatenated data according to Sramkó *et al.* (2019). Both maximum likelihood (ML) and Bayesian inference (BI) methods as implemented RAxML v.8.1.17 (Stamatakis 2014) and MrBayes v3.2.3 (Ronquist & Huelsenbeck 2003), respectively, were used for phylogenetic reconstruction.

The maximum likelihood (ML) analysis was performed with a GTR+G nucleotide substitution model, following the recommendations provided in the software manual. Branch support values were evaluated using 500 bootstrap replicates (MLBS) to assess topological robustness. For Bayesian inference (BI), the optimal substitution model was selected based on the Akaike information criterion (AIC) in jModelTest 2 (Darriba *et al.* 2012). The Bayesian analysis employed four independent Markov chain Monte Carlo (MCMC) runs, each initialized with random starting trees.

The chains were run for 2 million generations, with sampling every 1000th generation. After discarding the initial 20% of sampled trees as burn-in, a majority-rule consensus tree (>50%) was generated to estimate posterior probabilities (PP).

## Results and discussion

### Morphological affinity

The gross morphology of the newly found species is similar to *P. sukaczevii, P. tenuiloba* (Hayek 1904: 472) Juzepczuk (1937: 298), and *P. turczaninovii* Krylov & Sergievskaya (1930: 1), in that all of them have odd, finely lobed pinnate leaves. Among them, plants of *P. turczaninovii* are much larger than previous three and can be easily distinguished from them. *Pulsatilla tenuiloba* typically has leaves with more than five pairs of lateral leaflets, and the plant base usually bears numerous persistent withered petioles from previous years. The flowers are bluish-purple, and flowering occurs from June to late July. *Pulsatilla tenuiloba* is recorded in eastern Siberia, with its type locality being the Ingoda River (Juzepczuk 1937). It is very rare in China and is only found in northeastern Inner Mongolia (Zhao 1988). In contrast, the Hezijian population has leaves with four pairs of lateral leaflets, and the plants retain only a few withered petioles. Flowering begins in April, with flowers white inside and pale blue outside. By June, the fruits are already fully mature.

From herbarium materials, the Hezijian population is difficult to distinguish from *P. sukaczevii* There are great deal photos of Hezijian population online (Plant Photo Bank of China, PPBC. https://ppbc.iplant.cn/sp/14366) under the name of *P. sukaczevii. Pulsatilla sukaczevii* is recorded on the northern slope of Mount Krestovka, along the western shore of Lake Baikal (Juzepczuk 1937). It has also been documented in Inner Mongolia, China. A distinguishing feature of this species is its yellow or pale-yellowish flowers. Through field observations of wild *P. sukaczevii* populations in Ulanqab, Inner Mongolia, we confirmed that the Hezijian population clearly represents a distinct taxonomic entity from *P. sukaczevii*. Compared to the Hezijian population, *P. sukaczevii* from Ulanqab exhibits a more robust growth habit and typically inhabits drier, rocky or sandy slopes. Its flowers are bright-pale yellow to yellowish-white, with some individuals showing faint purple tinges at the base of the outer sepals. Notably, this species possesses a larger achene head and longer persistent styles than those observed in the Hezijian population (Figure 1). A detailed morphological comparisons between the Hezijian *Pulsatilla* and other similar species are provided in Table 1.

**TABLE 1.**
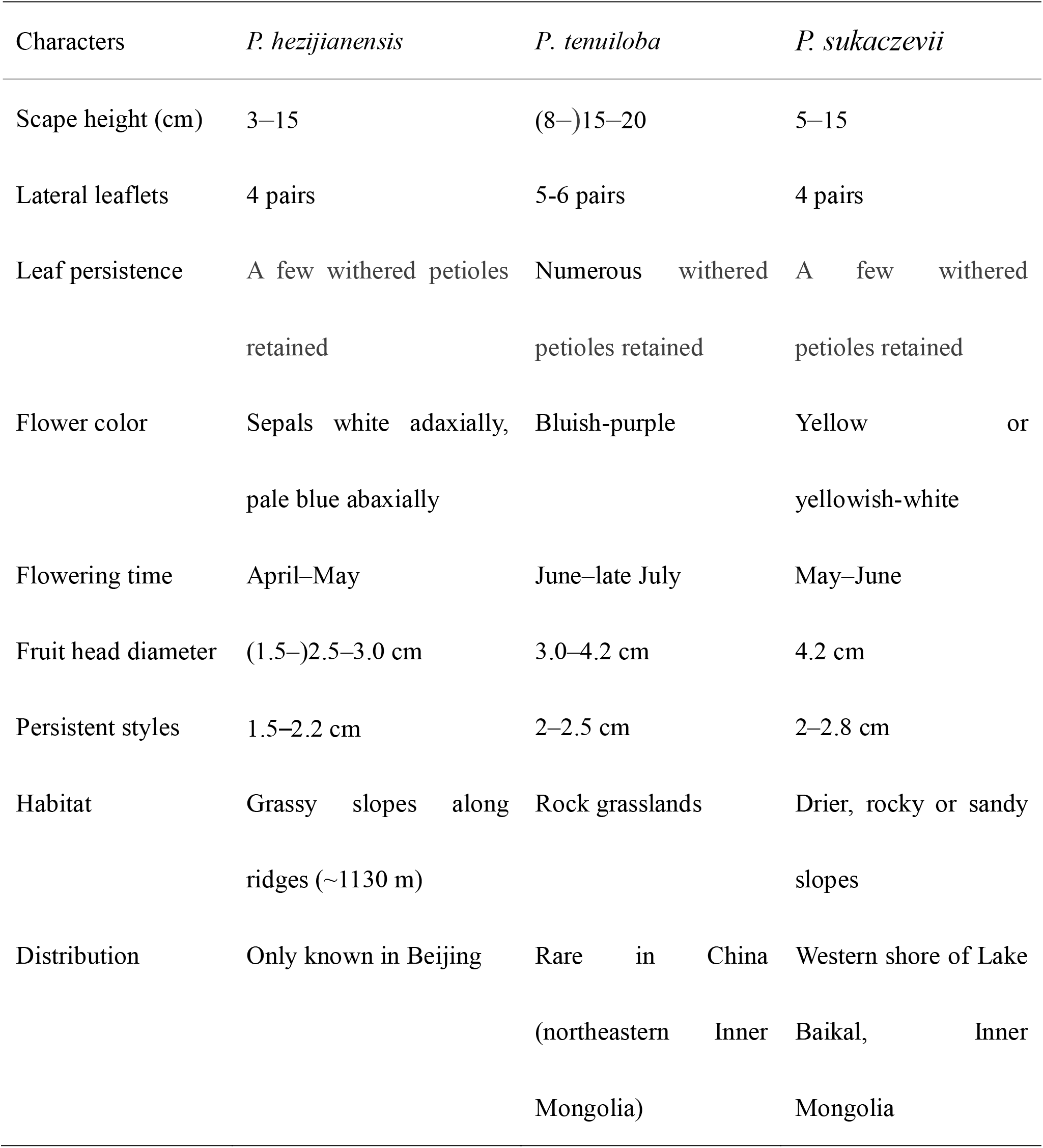
Morphological comparisons between *Pulsatilla hezijianensis, P. tenuiloba* and *P. sukaczevii*.

**FIGURE 1.**
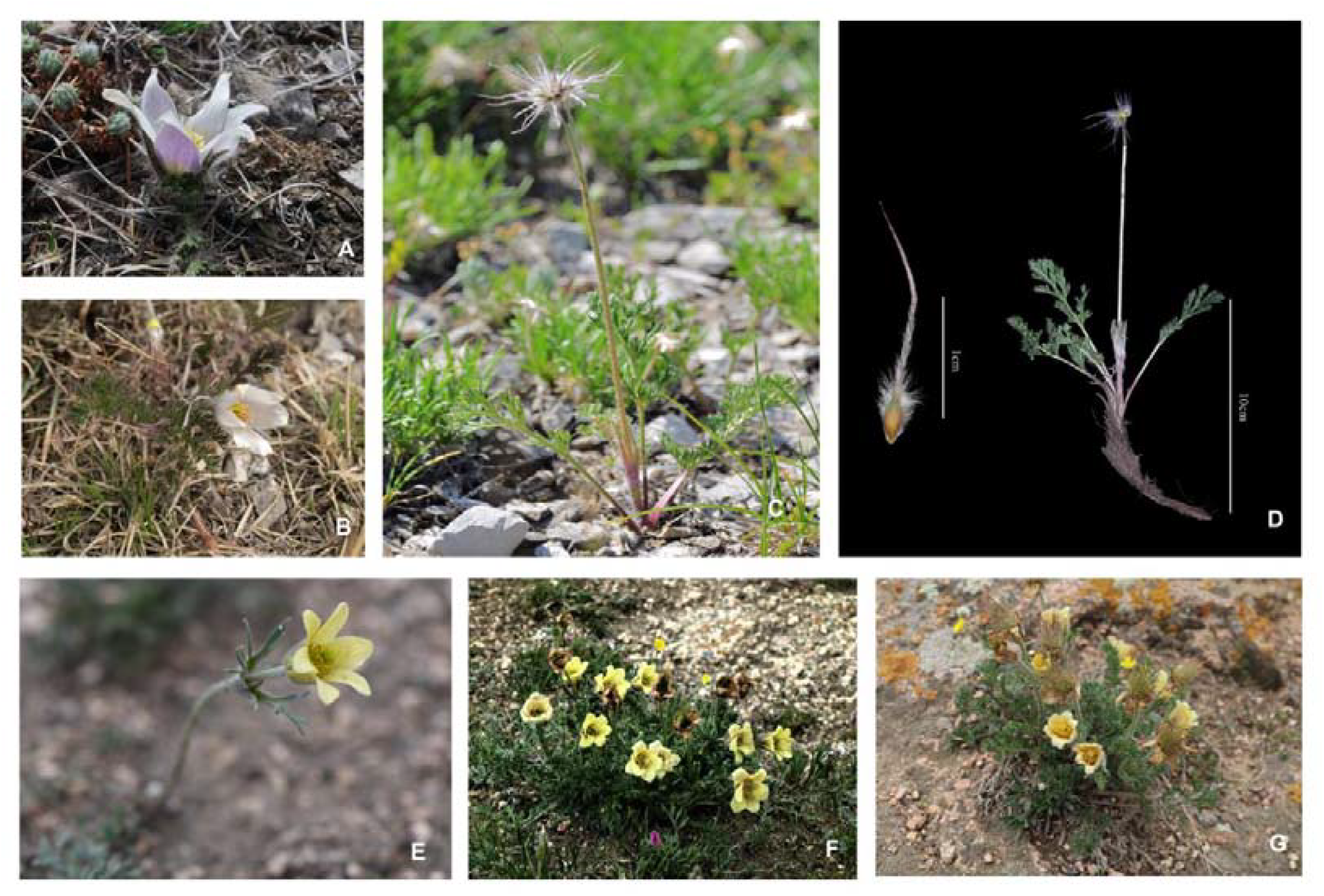
Photographs of *Pulsatilla hezijianensis* (A–D) and *P. sukaczevii* (E–G). A: Early flowering before leaf emergence; B: Leaves expanded by the late flowering stage; C: Fruit stage; D: Specimens illustrating achene size and scape height in fruit stage; E: An individual in flower; F–G: Dense clumps of *P. sukaczevii* in late flowering stage, and its more arid, rocky habitats.

### Phylogenetic position of the new species

The combined data matrix included a total of 2957 nucleotide bases, of which 233 (7.88%) were parsimony-informative sites. The major clades resolved with high support values were congruent between phylogenies reconstructed using Maximum Likelihood (ML) and Bayesian inference methods. Overall, the phylogenetic resolution within genus *Pulsatilla* remains limited, with several critical relationships among closely related species remaining unresolved (Figure 2). The phylogenetic framework and low phylogenetic resolution observed in *Pulsatilla* is consistent with previous Sanger sequencing-based phylogenetic reconstructions (Jiang *et al.* 2017, Sramkó *et al.* 2019, Xiang *et al.* 2025). The poor overall resolution might be caused by the lack of information in the limited gene regions used. Furthermore, *Pulsatilla* are known to readily hybridize between species (Grey-Wilson 2014) and it is likely to experience a recent rapid species radiation (Sramkó *et al.* 2019), making phylogenetic reconstruction difficult.

**FIGURE 2.**
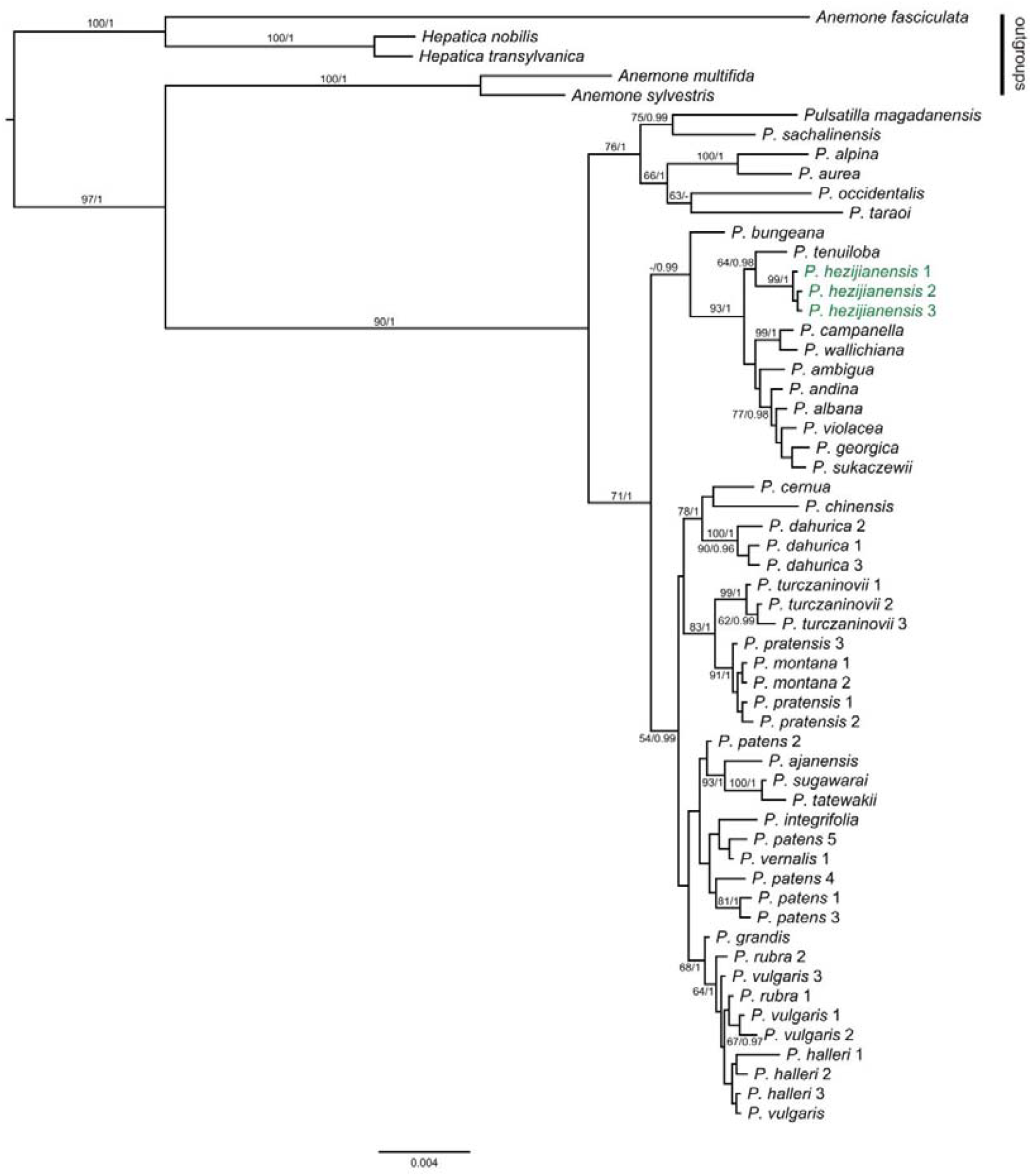
Bayesian phylogeny of trib. Ranunculeae obtained from the combined datasets. Numbers above and below branches are maximum likelihood bootstrap percentages (MLBS) and Bayesian posterior probabilities (PP), respectively. Only nodes with MLBS>50, and PP>0.95 were shown. Plant names in green are samples of new species, and those in black are retrieved from GenBank.

Despite of these difficulties, this study found that different individuals from the Hezijian population clustered together as a distinct clade and did not show close affinity to *P. sukaczevii*. We obtained consistent results from the nuclear gene phylogeny constructed using transcriptome data and coalescence methods, which also indicated no close relationship between Hezijian *Pulsatilla* with *P. sukaczevii*. However, due to limited taxon sampling of *Pulsatilla* in our phylogenomic analysis, these findings were not presented in this study. On the phylogenetic tree reconstructed in this study (Figure 2), the Hezijian population formed a sister relationship with *P. tenuiloba.* However, their substantial divergence was evident from the long branch lengths between them. Taking into account its aforementioned morphological differences, distinct distribution ranges, and flowering time divergence with *P. tenuiloba*, we propose that the *Pulsatilla* population from Hezijian village should be recognized as a new species. However, its origin, evolutionary history, and phylogenetic position require further clarification using phylogenomic data.

### Taxonomic treatment

**Pulsatilla hezijianensis** H. Y. Wu & L. Xie, *sp. nov.*, Fig. 1, A–D

#### Type

China, Beijing, Changping Dist., Hezijian, on grassy mountain ridges, 40.209041°N, 115.920389°E, alt. 1130 m, 26. Apr. 2023, *L. Xie et al., 2023042603* (holotype, BJFC). China, Beijing, Changping Dist., Hezijian, alt 1134 m, June 2025, *L. Xie et al., 2025062701* (BJFC).

#### Diagnosis

Morphologically allied to *P. tenuiloba* and *P. sukaczevii* but can be distinguished by its smaller and slender individuals (3–15 cm tall) especially in the late flowering and fruiting stages, shorter scapes, smaller fruit heads, sepals white adaxially and pale blue abaxially.

#### Etymology

The specific epithet refers to the type locality, Hezijian village, Changping Dist. Beijing.

#### Description

Herbs perennial, small, 3–15 cm tall. Rhizome vertical, ca. 5–8 cm, robust. Leaves all basal, 4–6, not expanded at anthesis; petiole ca. 3 cm, villose; leaf blade ovate to narrowly ovate to oblong-ovate, 3–4 × 1.5–2.2 cm, with 4 pairs of lateral leaflets, 2 or 3 × pinnately divided, villose at tip of the terminal lobes, apex acute, terminal lobes lanceolate, 0.5–0.8 mm wide. Scape 1, erect, 1.5–2.5 cm, elongated to 12 cm in fruit, villose; involucral bracts ca. 1.0 cm, densely villose, basally connate into a 2–3 mm tube, apically finely divided. Sepals white inside, white to pale-blue outside, erect or ascending, oblong-ovate, 1–2 × 0.4–1 cm, abaxially thickly puberulent, apex slightly acute to round. Anthers yellow. Infructescences ca. (1.5–)2.5–3.0 cm in diam. Achenes 2.5–3 mm, densely puberulent. Persistent styles 1.5–2.2 cm, basally puberulent, apically pilose.

### Phenology

Flowering in April to May, and fruiting from May to June.

### Vernacular name

The Chinese mandarin name is “Hé Zi Jiàn Bái Tóu Wēng” (禾子涧白头翁)

#### Habitat

Grassy slopes along ridges in montane areas; ca. 1130 m.

#### Ecology, distribution and status

Endemic to Beijing. This species was known only from its type locality. In Hezijian village, Changping Co., Beijing, individuals of this species grow on grassy slopes along the mountain ridge. The area we discovered is rather small, including a few dozen individuals. This species’s habitat is highly vulnerable to anthropogenic disturbances, which may lead to population degradation or even local extinction. According to the IUCN red list categories and criteria (IUCN Standard and Petitions Committee 2022), *Pulsatilla hezijianensis* should be categorized as critically endangered (CR).

## Acknowledgments

This research was funded by the National Natural Science Foundation of China (grant number: 31670207).

---

**TABLE S1**.Species, GenBank accession numbers, voucher information of the sequences used in this study.

## References

Aichele, D. & Schwegleb, H.W. (1957) Die Taxonomie der Gattung Pulsatilla. Repertorium Novarum Specierum Regni Vegetabilis 60: 1–230.

Bunge, A. A. (1833) Enumeratio plantarum, quas in China boreali collegit Dr. Al.

Bunge. Anno 1831. Mémoires Savantes Etranges Académie Sciences St.-Petersbourg 2: 75–147.

Darriba, D., Taboada, G.L., Doallo, R. & Posada, D. (2012) jModelTest 2: More models, new heuristics and parallel computing. Nature Methods 9(8): 772. 10.1038/nmeth.2109

Grey-Wilson, C. (2014) Pasque-flowers. The genus Pulsatilla. The Charlotte-Louise Press, Kenninghall.

Hayek, A. (1904): Kritische Übersicht über die Anemone-Arten aus der Section Campanaria Endl. und Studien über deren phylogenetischen Zusammenhang. In: Urban, I. & Graebner, P. (Eds.): Festschrift zur Feier des Siebzigsten Geburtstages des Herrn Professor Dr. Paul Ascherson. Gebr. Bornträger, Leipzig, pp.451–475..

Hoot, S.B., Meyer, K.M. & Manning, J.C. (2012) Phylogeny and reclassification of Anemone (Ranunculaceae), with an emphasis on austral species. Systematic Botany 37(1): 139–152. 10.1600/036364412X616738

IUCN Standards and Petitions Committee. 2022. Guidelines for Using the IUCN Red List Categories and Criteria. Version 15. Gland: IUCN.

Jiang, N., Zhou, Z., Yang, J.B., Zhang, S.D., Guan, K.Y., Tan, Y.H. & Yu, W.B. (2017) Phylogenetic reassessment of tribe Anemoneae (Ranunculaceae): Non-monophyly of Anemone s.l. revealed by plastid datasets. PLoS ONE 12(3): e0174792. 10.1371/journal.pone.0174792

Juzepczuk, S.V. (1937) Pulsatilla Adans. In: Komarov, V.L. & Schischkin, B.K. (Eds.) Flora of the USSR. Vol. 7. Editio Academiae Scientiarum URSS, Moscow & Leningrad, pp. 285–307.

Katoh, K., Kuma, K.I., Toh, H. & Miyata, T. (2005) MAFFT version 5: Improvement in accuracy of multiple sequence alignment. Nucleic Acids Research 33: 511–518. 10.1093/nar/gki198

Kearse, M., Moir, R., Wilson, A., Stones-Havas, S., Cheung, M., Sturrock, S., Buxton, S., Cooper, A., Markowitz, S., Duran, C., Thierer, T., Ashton, B., Meintjes, P. & Drummond, A. (2012) Geneious Basic: an integrated and extendable desktop software platform for the organization and analysis of sequence data. Bioinformatics 28(12): 1647–1649. 10.1093/bioinformatics/bts199

Krylov, P. N. & Sergievskaya, L. P. (1930) Pulsatilla turczaninovii sp. n. Sistematicheskie Zametki po Materialam Gerbariya imeni P. N. Krylova Tomskogo Gosudarstvennom Univiversiteta imeni V. V. Kuybysheva 1930(5–6): 1–2.

Li, Q.J., Su, N., Zhang, L., Tong, R.C., Zhang, X.H., Wang, J.R., Chang, Z.Y., Zhao, L. & Potter, D. (2020) Chloroplast genomes elucidate diversity, phylogeny, and taxonomy of Pulsatilla (Ranunculaceae). Scientific Reports 10(1): 19781. 10.1038/s41598-020-76699-7

Li, Q.J., Wang, X., Wang, J.R., Su, N., Zhang, L., Ma, Y.P., Chang, Z.Y., Zhao, L. & Potter, D. (2019) Efficient identification of Pulsatilla (Ranunculaceae) using DNA barcodes and micro-morphological characters. Frontiers in Plant Science 10: 1196. 10.3389/fpls.2019.01196

Linnaeus, C. (1753) Species plantarum, exhibentes plantas rite cognitas ad genera relatas, cum differentiis specificis, nominibus trivialibus, synonymis selectis, locis natalibus, secundum systema sexuale digestas. Vol. 1. Laurentii Salvii, Holmiae, 538 pp. 10.5962/bhl.title.669

Miller, P. (1754) The Gardener’s Dictionary, ed. 4. Privately published, London (unpaged).

Regel, E.A. (1861) Tentamen florae ussuriensis oder Versuch einer Flora des Ussuri-Gebietes nach den von Herrn R. Maack gesammelten Pflanzen bearbeitet. Mémoires de l’Académie Impériale des Sciences de Saint Pétersbourg, sér. 7, 4 (4): 1–228.

Ronikier, M., Costa, A., Aguilar, J.F., Feliner, G.N., Küpfer, P. & Mirek, Z. (2008) Phylogeography of Pulsatilla vernalis (L.) Mill. (Ranunculaceae): chloroplast DNA reveals two evolutionary lineages across central Europe and Scandinavia. Journal of Biogeography 35(9): 1650–1664. 10.1111/j.1365-2699.2008.01907.x

Ronquist, F. & Huelsenbeck, J.P. (2003) MrBayes 3: Bayesian phylogenetic inference under mixed models. Bioinformatics 19(12): 1572–1574. 10.1093/bioinformatics/btg180

Sramkó, G., Laczkó, L., Volkova, P.A., Bateman, R.M. & Mlinarec, J. (2019) Evolutionary history of the Pasque-flowers (Pulsatilla, Ranunculaceae): Molecular phylogenetics, systematics and rDNA evolution. Molecular Phylogenetics and Evolution 135: 45–61.

Stamatakis, A. (2014) RAxML version 8: A tool for phylogenetic analysis and post-analysis of large phylogenies. Bioinformatics 30(9): 1312–1313. 10.1093/bioinformatics/btu033

Tamura, M. (1995) Trib. Ranunculeae. In: Hiepko, P. (Ed.) Die Natürlichen Pflanzenfamilien, vol. 17a. Duncker & Humbolt, Berlin, pp. 356–366.

Thiers, B. (2017) Index Herbariorum. A global directory of public herbaria and associated staff. New York Botanical Garden’s Virtual Herbarium.

Wang, W.T. & Bartholomew, B. (2001) Ranunculaceae (36): Pulsatilla. In: Wu, Z.Y. & Raven, P.H. (Eds.) Flora of China, vol. 6. Science Press, Beijing & Missouri Botanical Garden Press, St. Louis, pp. 329–333.

Xiang, K.-L., Cao, G.-L., Peng, H.-W., Li, X.-Q., Erst, A.S., Lian, L., Erst, T.V., Jabbour, F., Ortiz, R. del C. & Wang, W. (2025) Molecular phylogeny of Anemone (Ranunculaceae), with special emphasis on the origin of alpine species. Taxon 74(2): 292–306. 10.1002/tax.13297

Zhang, T.-T., Zhang, S.-M., Xu, L. & Kang, T.-G. (2022) Pulsatilla saxatilis (Ranunculaceae), a new species from north-east China. Phytotaxa 539(2): 195–202. 10.11646/PHYTOTAXA.539.2.6

Zhao, Y.-Z. (1988) Classification and eco-geographical distribution of the genus Pulsatilla Adans. in Nei Mongol. Acta Scientiarum Naturalium Universitatis Intramongolicae 19(4): 654–661.

